# Excited-state dynamics of the all-trans protonated retinal Schiff base in CRABPII-based rhodopsin mimics

**DOI:** 10.1101/2022.02.16.480777

**Authors:** Gaoshang Li, Yongnan Hu, Sizhu Pei, Jiajia Meng, Jiayu Wang, Ju Wang, Shuai Yue, Zhuan Wang, Shufeng Wang, Xinfeng Liu, Yuxiang Weng, Xubiao Peng, Qing Zhao

**Author notes:** Draft manuscript. Please do not cite without the authors’ permission.

## Abstract

The rhodopsin mimic is a chemically synthetized complex with the retinyl Schiff base (RSB) formed between protein and the retinal chromophore, which can mimic the natural rhodopsin-like protein. The artificial rhodopsin mimic is more stable and designable than the natural protein and hence has wider uses in photon detection devices. The mimic structure RSB, like the case in the actual rhodopsin-like protein, undergoes isomerization and protonation throughout the photoreaction process. As a result, understanding the dynamics of the RSB in the photoreaction process is critical. In this study, the transient absorption spectra (TAS) of three mutants of the cellular retinoic acid-binding protein II (CRABPII)-based rhodopsin mimic at PH = 3 were recorded, from which the related excited-state dynamics of the all-trans protonated RSB (AT-PRSB) were investigated. The transient fluorescence spectra (TFS) measurements are used to validate some of the dynamic features. We find that the excited-state dynamics of AT-PRSB in three mutants share a similar pattern that differs significantly from the dynamics of 15-cis PRSB (15C-PRSB) of the rhodopsin mimic in neutral solution. By comparing the dynamics across the three mutants, we discovered that the aromatic residues near the β-ionone ring structure of the retinal may help stabilize the AT-PRSB and hence slow down its isomerization rate. Furthermore, from the three mutants, we find one protein with near-infrared fluorescence emission up to 688 nm, leading to further possible applications in sensing or bioimaging.

## Introduction

The quantum effect in biology has been a hot topic that has attracted attentions in the last few decades(1; 2; 3). One of the most famous examples is the photoreaction process in photosensitive proteins such as rhodopsin(4), bacteriorhodopsin(BR)(5), proteorhodopsin (PR)(6), and Anabaena sensory rhodopsin (ASR)(7), *etc*. Rhodopsin-like proteins are widely spread in nature and have similar structures. They are typically made up of a seven-helical transmembrane protein and a chromophore named retinal, which can switch its conformations when absorbing photons. Although there are many distinct isomers of retinal in solution, the isomerization of the retinal in rhodopsin is almost unique. In bacteria and microbes, the isomerization is 13-trans/cis, allowing the rhodopsin-like proteins to perform the biological functions such as light-driven proton transport, chloride pumping or light sensing; In vertebrate animals the isomerization is 11-cis/trans, resulting in the signal transduction for vision. In the literature (8) the authors give a systematic review on the rhodopsin-like proteins. Although the mechanisms and the roles of the rhodopsin-like proteins differ significantly, they are all sensitive to the dim light. Understanding the mechanisms of their photoreaction process can help in the construction of a novel photon detecting device with great sensitivity. The rhodopsin-like protein has been a representative of the photosensitive proteins. On the mechanism of different rhodopsin-like proteins, we refer to the literature(9), where we can see that the photoreaction dynamics depend on the type of the rhodopsin-like protein and its environment, such as the pH value.

Natural rhodopsin-like proteins usually contain more than 300 residues and are embedded in the membrane. As a result, the experimental measurements and the computational simulations on the natural rhodopsin-like proteins are complicated and costly. Fortunately, the synthetical protein can provide a low-cost and simple investigation on the mechanism of photoreaction in rhodopsin-like protein from two aspects. On one hand, the rational design allows the synthetical protein to be much smaller and independent on the membrane while retaining the major photoreaction capabilities. On the other hand, the designable proteins are relatively stable on limited mutations, allowing us to study the photoreaction mechanisms through systematical mutagenesis. Geiger, Borhan *et al*.have been working on the artificial rhodopsin mimics for a long history and have provided several mimics with the human proteins CRABPII or the cellular retinol binding protein II (CRBPII) as scaffoldings (10; 11; 12; 13). The crucial component for the photoreaction, the retinal Schiff base (RSB), is correctly generated in these mimics between the chromophore retinal and the sidechain of the residue LYS in the scaffolding protein. They also investigated the spectral characteristics and photoreaction dynamics of the RSB in several of the rhodopsin mimics. They tuned the light absorption in a wide spectral range from purple to red using point mutations around the RSB region in CRABPII-based mimic (10) and CRBPII-based mimic (14); They explored the dynamics of the AT-PRSB in CRBPII-based rhodopsin mimics from both TAS experiments and simulations (15; 16); They also revealed the conversion between 15C-PRSB and all-trans unprotonated RSB (AT-URSB) in a Hexa-mutant (M2-KFVQ:P39Y: R59Y) of the CRABPII-based rhodopsin mimics in the neutral solution (17). However, the dynamics of the AT-PRSB in the CRABPII-based rhodopsin mimics have not yet been investigated.

This study tries to solve this question by evaluating both the TAS and the TFS of PRSB in three mutants of the CRABPII-based rhodopsin mimics in acidic solution. From the available mutants listed in literature (11), we select three known as M1(KFVQ:P39Q:R59Y), M2(KFVQ:P39Y:R59Y) and M9 (KFVQ:P39Y:R59Y:A32W) as the subjects of our study, where from M1 to M9 more non-aromatic residues are mutated to aromatic ones. We discover a general pattern for the photoreaction process of the AT-PRSB in the CRABPII-based rhodopsin mimic in acidic solution, by comparing the excited-states dynamics across the three chosen mutants. The different mutations in CRABPII-based mimic only affect the detailed absorption/fluorescence spectral properties and the reaction rates. By further relating the spectra to the structural properties of the three mutants, we infer that the aromatic residues near the *β*-ionone ring structure can stabilize the PRSB and slow down the isomerization process. Furthermore, when we compare the fluorescence properties of the PRSB across the three mutations, we discover that mutant M9 exhibits the brightest fluorescence and the longest wavelength among the three mimics. In particular, the fluorescence of mutant M9 is in the far red region, which may be useful in molecular devices design such as the sensors or microscopes for bioimaging in deeper tissues(18).

## Results

The structures of the three mutants are shown in Fig 1, with mutants M1 and M2 being the crystallographic structures directly taken from the Protein Data Bank (PDB) (11). Since there is no available structure for mutant M9 in PDB, we give its representative structure with simple molecular dynamics (MD) simulations. In detail, we first generate the initial structure of mutant M9 by point mutation from M2 using the UCSF Chimera software (19) with the Dunbrack 2010 rotamer library (20) plug-in, then perform the plain MD simulation for 200ns at 310K in explicit solvent and finally, select the most representative structure from the trajectory. As indicated in Fig 1, the overall structures of the three mutants are very similar, except the local environment nearby the *β*-ionone ring of the chromophore retinal. With the three mutants in PBS solution, we first measure their steady absorption and fluorescence spectra, followed by the time-resolved spectral known as the transient absorption spectra (TAS) and transient fluorescence spectra (TFS), from which the excited-state dynamics after photoactivation are analyzed using the global analysis techniques (21).

**Figure 1:**
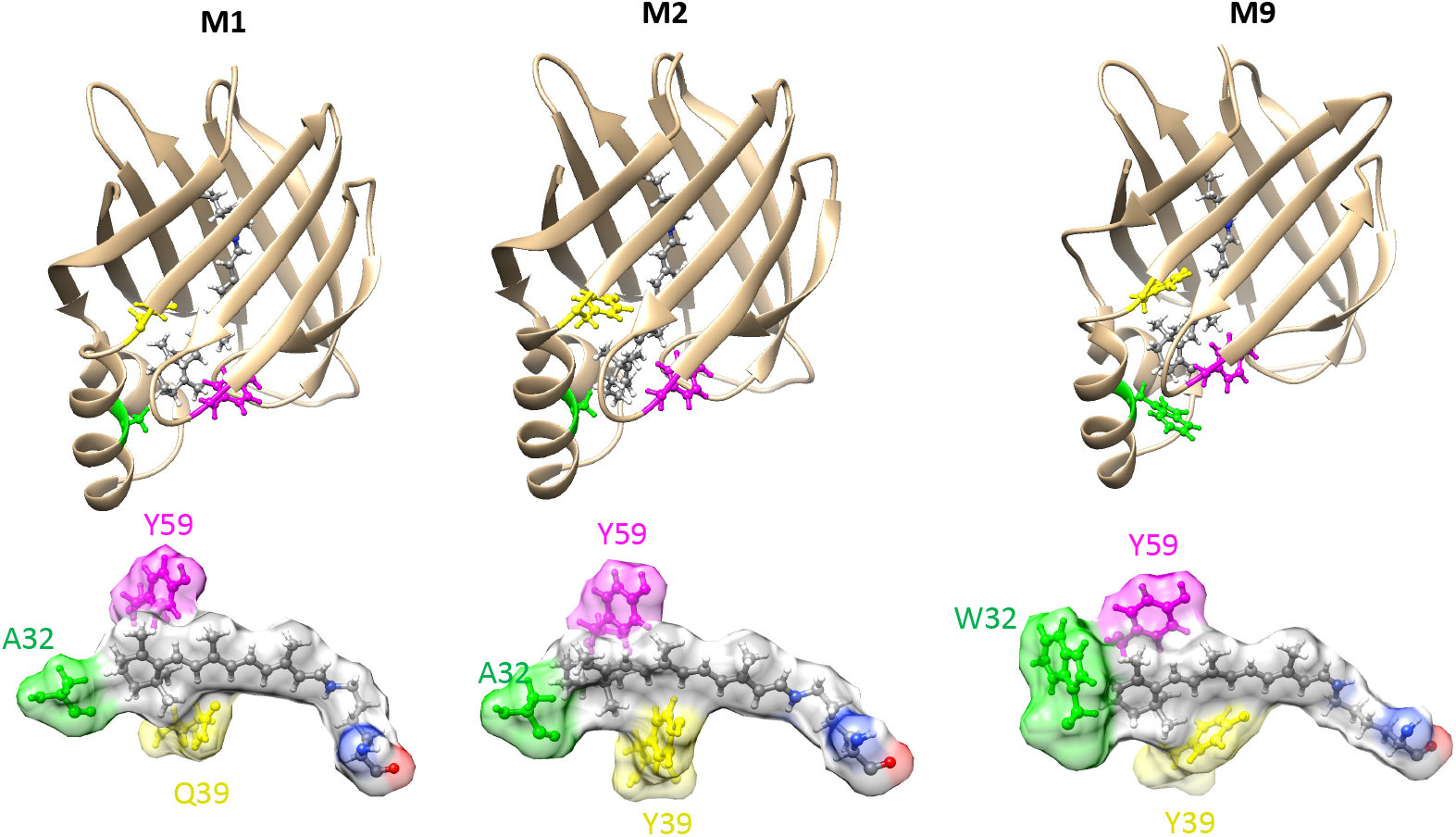
The structure of the mutants M1, M2 and M9. The bottom row shows the zoomed-in local environment nearby the retinal chromophore for each mutant.

### Steady absorption and fluorescence spectra

In Fig 2A and Fig 2B, we show the normalized UV–vis absorption spectra and fluorescence spectra for three mutants, and we can see that the three mutants have similar lineshapes for both. We notice that the mutants M1 and M2 are well distinguished from the steady absorption spectra but cannot be distinguished from the steady fluorescence spectra.

**Figure 2:**
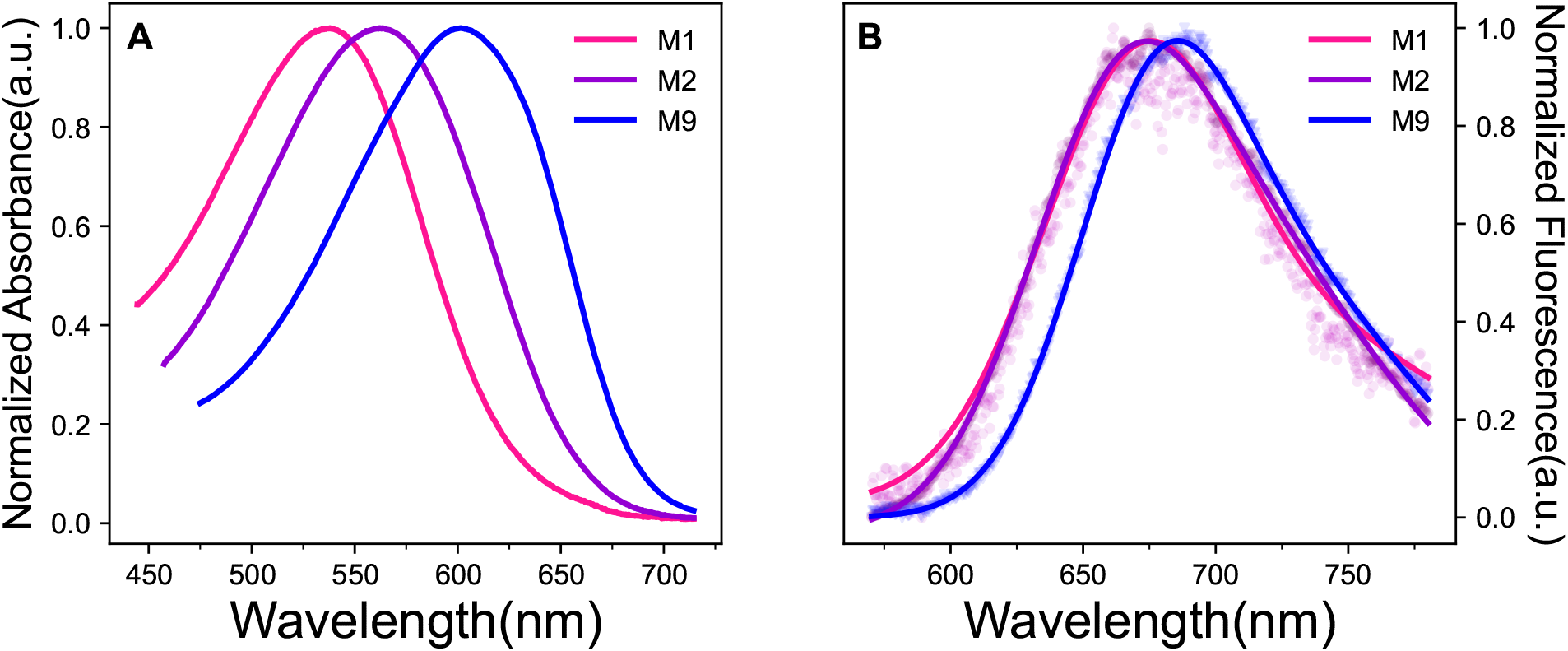
The normalized steady absorption(A) and fluorescence spectra for mutants(B) M1, M2 and M9.

The wavelengths of maximal absorbance and emission for each mutant are identified from Fig 2 and are listed in Table 1, where we can see that all exhibit substantial Stokes Shifts with more than 80 nm. We notice that our measurements on the wavelength of maximum absorbance for three mutants are well consistent with the values of PRSB in low-pKa states as listed in reference (11), indicating that all three mutants in our system are initially in the dark state, i.e., the AT-PRSB state.

**Table 1:**
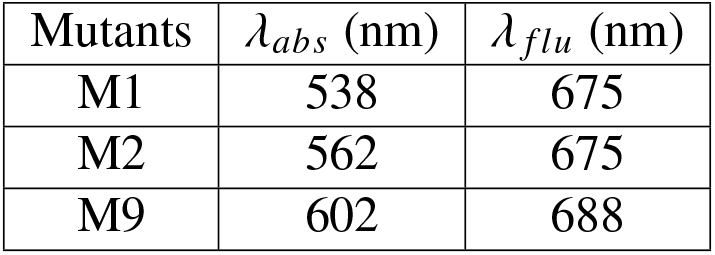
The wavelengths of maximum absorption and fluorescence spectra for three mutants

| Mutants | $\lambda_{abs}$ (nm) | $\lambda_{flu}$ (nm) |
| --- | --- | --- |
| M1 | 538 | 675 |
| M2 | 562 | 675 |
| M9 | 602 | 688 |

By comparing the wavelength of maximum absorption *λ_abs_* and the wavelength of maximum emission *λ_flu_* across three mutants, we have the relations *λ*_*abs,M*1_ < *λ*_*abs,M*2_ < *λ*_*abs,M*9_ and *λ*_*flu,M*1_ = *λ*_*flu,M*2_ < *λ*_*flu,M*9_. In particular, mutant M9 has the fluorescence wavelength in far red region (688nm) as well as the strongest brightness, so it has great potential to be applied in sensing or bioimaging. Recalling the local environment nearby retinal as depicted in Fig 1, we notice that the relation of *λ_abs_* is well correlated with the number of the aromatic sidechains nearby the *β*-ionone ring structure of the retinal in the mutants. With more aromatic sidechain nearby the *β*-ionone ring of the retinal, the absorption spectra becomes more red-shifted. We infer that there is stronger *π* stacking interaction with more aromatic sidechains nearby the *β*-ionone ring, which can affect the distributions of the electron along the retinal. In the following sections, we will further investigate the influence of the aromatic sidechains on the excited-state dynamics of the PRSB.

### The transient absorption spectra (TAS)

We perform the ultrafast TAS measurements on different mutants with the 560nm pump and white-light probe pulses, covering the time scale from femtoseconds to nanoseconds. The color-coded contour maps of the changes in optical density as a function of time and probe wavelength (Δ*OD*(*λ, t*)) are shown in Fig S1. In Fig 3 we show the absorption spectra at several selected time points for mutants M1 (left), M2 (middle) and M9 (right), respectively. We can see that the three mutants have similar patterns in TAS. In the TAS of each mutant, there are three obvious peaks corresponding to the excited state absorption (ESA), ground state bleaching (GSB) and stimulated emission (SE), respectively. The positive peaks are all located at around 470nm, indicating that all the three mutants share similar energy level structures for the ESA. However, their decay times are different, showing that the mutants have different lifetimes for the excited-states, where M1 has the shortest lifetime while M9 has the longest lifetime. The negative peaks due to GSB are located at around 560nm, 570nm and 600nm for mutants M1, M2 and M9, respectively, consistent with their steady absorption spectra. Furthermore, in wavelength interval 700–750nm for each mutant, there is significant but relatively weak negative signal different from the GSB signal, indicating that the SE process happens there, which we will further validate using the TFS in the next section. The peaks for SE are located at around 710nm, 701nm and 690nm for mutants M1, M2 and M9, respectively. For each mutant, the wavelength at the peak of SE is clearly longer than that in the steady fluorescence spectra, which can be *partly* explained by the relation between the SE spectra and the spontaneous emission (fluorescence) spectra as revealed in reference (22). In the end, we note that all the three mutants exhibit a slight red-shifting of the SE and a clear blue-shifting of the ESA (see the dashed lines in Fig 3) with time going on, which is attributed to the vibrational relaxation on the excited-states potential energy surface (PES) from the initial Frank-Condon (FC) state.

**Figure 3:**
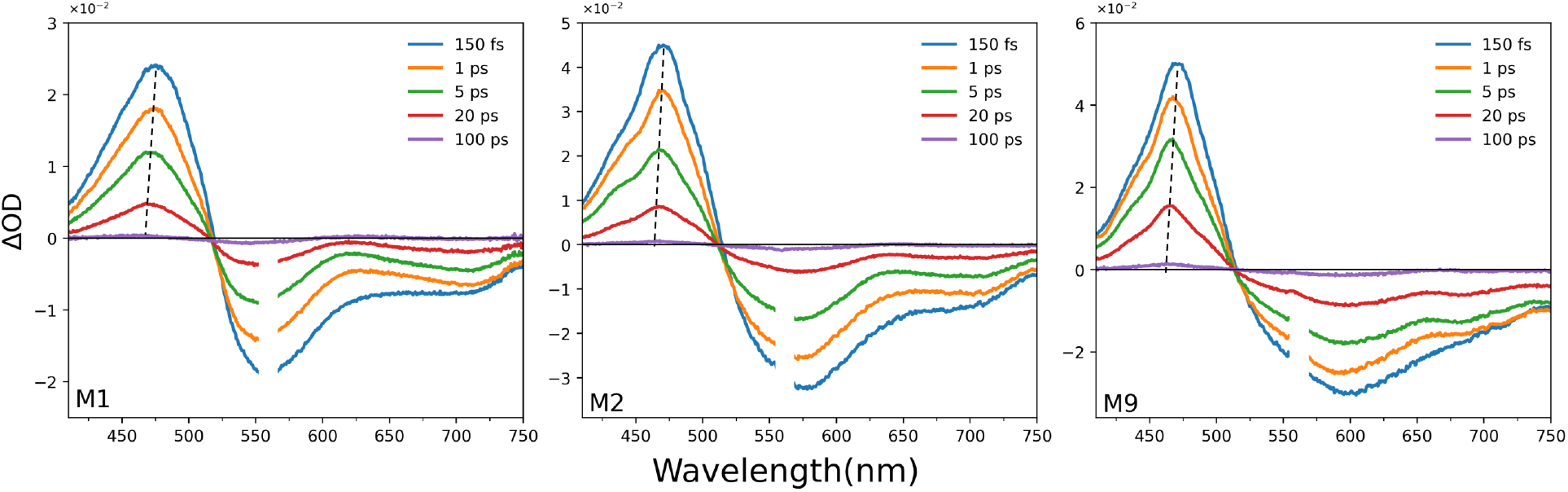
The absorption spectra at different time points for mutants M1(left), M2(middle) and M9(right).The 560 nm region is disregarded due to the low signal-to-noise ratio by pump scattering.

By performing the global analysis on the TAS, we give the model of the photoreaction process for each mutant. We find that the TAS data can be well fitted using a sequential model with four compartments. The evolution-associated difference spectra (EADS) and the concentration profiles for the three mutants are shown in Fig 4.

**Figure 4:**
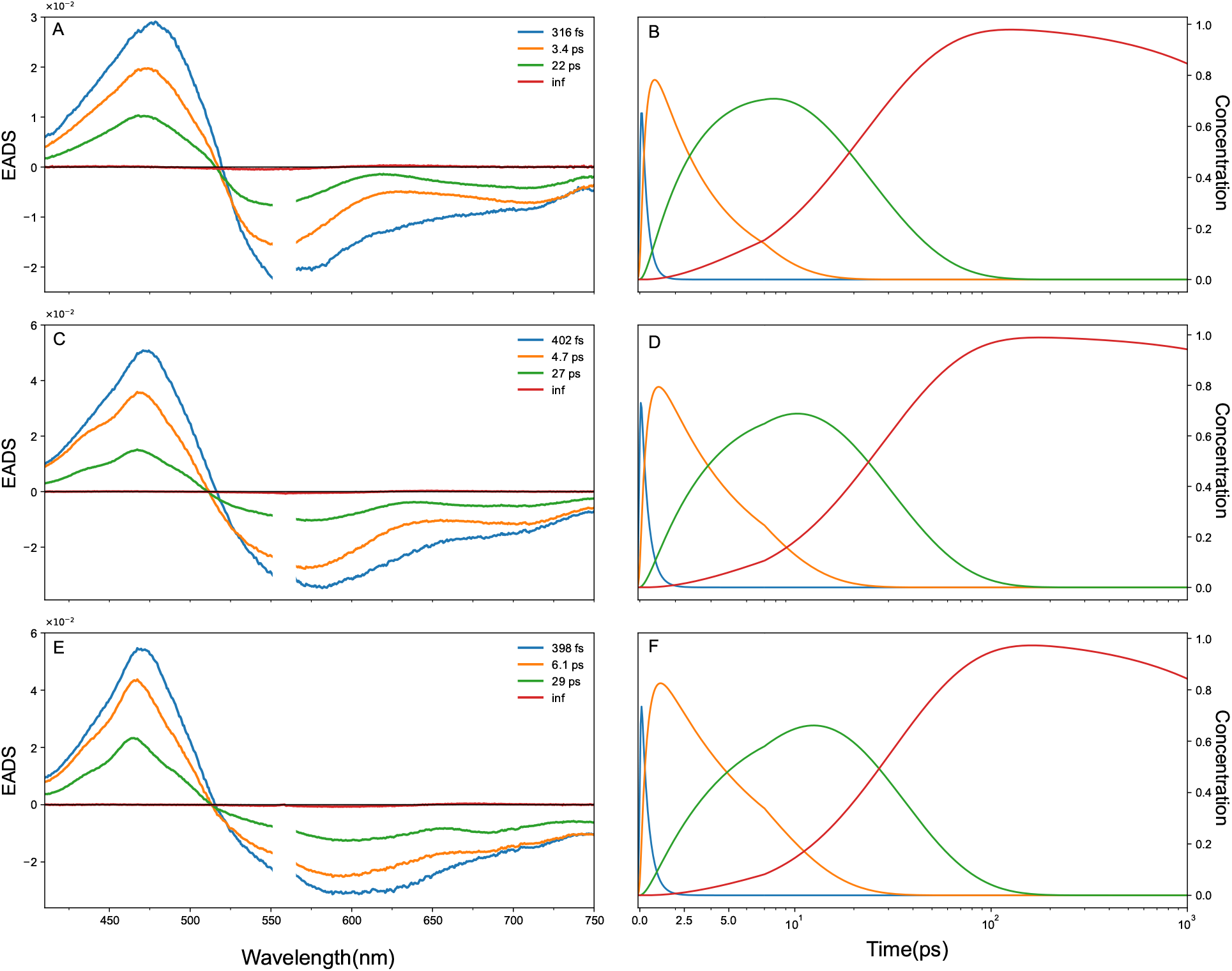
The EADS obtained from the global analysis on the TAS.

Taking the mutant M2 as an example, by global analysis we obtain four compartments with lifetime 402fs, 4.7ps, 27ps and 16ns, with their corresponding EADS as shown in Fig 4B. Since the obtained lifetime for the fourth compartment is much larger than our experimental time scale, we take it as infinity. We interpret the sequential model with four compartments as shown in Fig 5. The system is firstly excited to the FC state by the pulse at 560nm, and then decays to the nearest local minimum on the excited-state PES within 440fs. From such a local minimum the system overcomes a low energy barrier and reaches a second local minimum in 5ps. Afterwards, the system continues to overcome an even higher energy barrier and reach the canonical intersection (CI) point, from which it isomerizes to the ground state of the 15C-PRSB, taking about 27ps in total. In the end, the ground state of 15C-PRSB takes a very long time to relax back to the ground state of the AT-PRSB, corresponding to the fourth compartment with infinite lifetime. The EADS of the fourth compartments is too weak to be seen compared to the other three in Fig 4B, so we show the zoomed-in spectrum of the fourth compartment in Fig S2B. In addition to the GSB peak at around 560 nm, we found another positive peak at 650 nm for the fourth compartment, which was never observed previously. We attribute this peak to the ESA of 15C-PRSB in the acidic solution.

**Figure 5:**
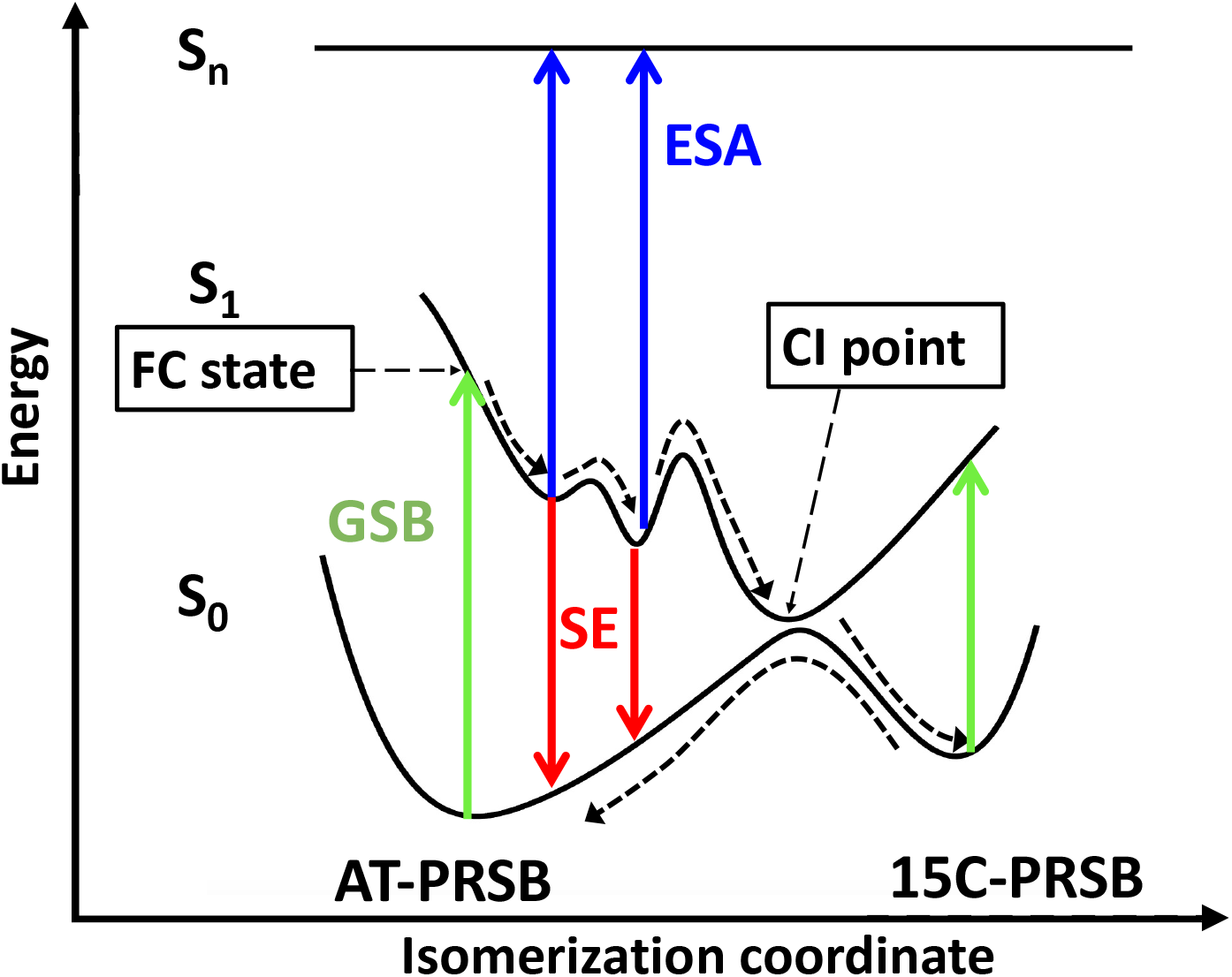
The proposed interpretation on the four-compartment sequential model on TAS of the AT-PRSB in CRABPII-based rhodopsin mimic in acidic solution. The corresponding photoreaction processes including the GSB, ESA and SE are labeled in different colors corresponding to their respective wavelengths.

We get similar conclusions on the excited-state dynamics for the other two mutants M1 and M9 in Fig 4A and Fig 4C, respectively. The main difference among three mutants lies in the lifetime of the compartments and the details of the spectra. In Table 2, we list the lifetime values of the four compartments obtained from global analysis for different mutants. In particular, we find that the lifetimes of the second and third compartments both increase in the order of M1, M2 and M9, which are again consistent with the increase in the number of the aromatic sidechain nearby the *β*-ionone ring structure of retinal from M1 to M9. Since the lifetime of the third compartment is corresponding to the isomerization process, we hereby infer that the aromatic sidechain nearby the *β*-ionone ring structure of the retinal can effectively increase the retinal’s stability, making the isomerization difficult. Similar to the property of the steady absorption spectra, the nearby aromatic sidechain also leads to a red-shift in the wavelength of GSB in the mutant M9 compared to M1 and M2.

**Table 2:** The lifetimes of the four compartments obtained from global analysis for different mutants.

| Mutants | $\tau_1$ (fs) | $\tau_2$ (ps) | $\tau_3$ (ps) | $\tau_4$ (ps) |
| --- | --- | --- | --- | --- |
| M1 | 316 | 3.4 | 22 | inf |
| M2 | 402 | 4.7 | 27 | inf |
| M9 | 398 | 6.1 | 29 | inf |

### The transient fluorescence spectra (TFS)

To further validate our analysis on the excited-state dynamics, we measure the TFS of the AT-PRSB in the mutants M1, M2, M9 as show in Fig S3, where the mutants M1 and M9 have the weakest and strongest fluorescence, respectively. From Fig S3, we can see that the wavelengths at the peaks of the TAS are well consistent with the values in the steady fluorescence spectra as listed in Table 1. With a single exponential model, we can fit the TFS well. In Fig 6 we show the fitting results on the decay curves at the peak of the fluorescence spectra for each mutant. The lifetime values of the fluorescence, which are usually also the lifetimes for the excited states, are obtained as 30ps, 33ps and 34ps for M1, M2 and M9, respectively. Recalling the results of the global analysis on the TAS as shown in Table 2, the lifetime of the excited states is given by *τ* = *τ*_1_ + *τ*_2_ + *τ*_3_ for the sequential model, with *τ*_1_, *τ*_2_, *τ*_3_ are the lifetimes for the first three compartments. Accordingly, the lifetimes of the excited state for three mutants given by the TAS analysis are 26ps, 32ps and 35ps, respectively. We can see that the lifetimes of the excited states obtained from the TFS fitting are well consistent with the values from the global analysis on the TAS. There is relatively larger difference in the lifetime of excited states between the TFS and TAS analysis for mutant M1, which we attribute to the uncertainties in the experimental measurements due to the weak fluorescence of mutant M1.

**Figure 6:**
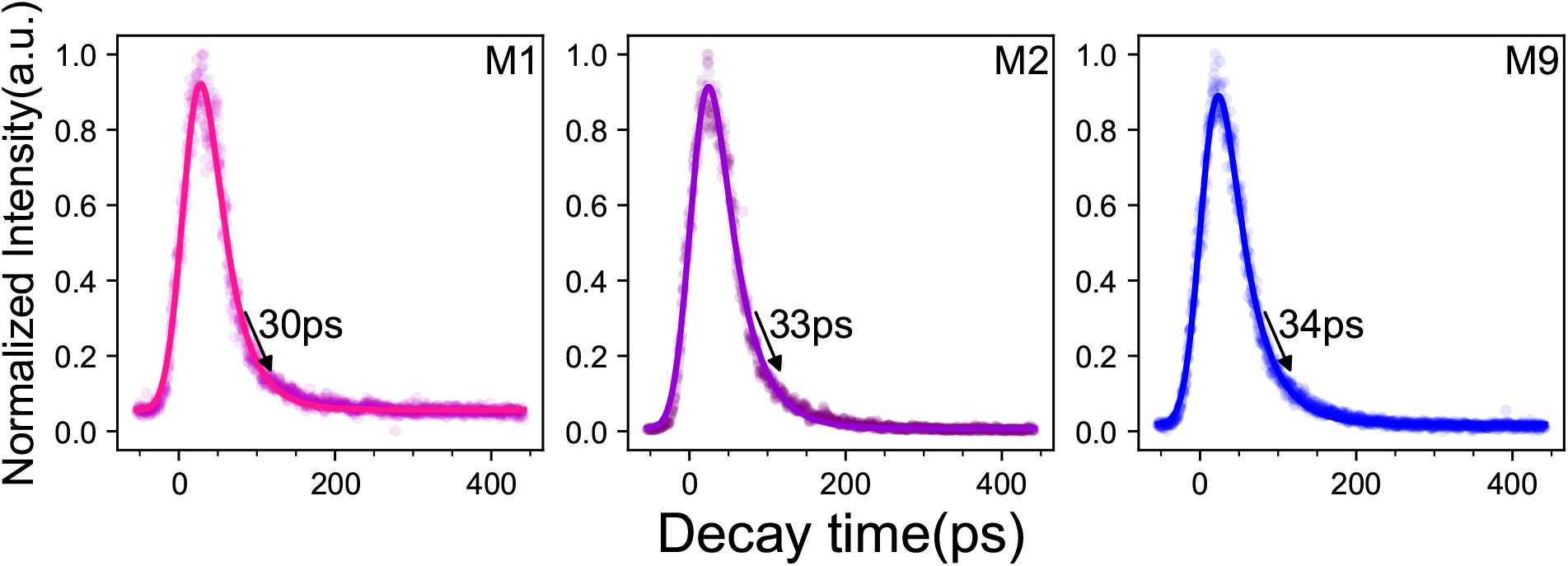
Fitting results on the decay curves at the peak of the fluorescence spectra for three mutants with a single exponential model.

In the end, we compare the properties of the TFS across the three mutants. We find that mutant M9 has the brightest fluorescence, longest wavelength and longest lifetime compared with the other two, leading to wider applications in biosensing or bioimaging. Again, such an observation is consistent with the fact that M9 has the largest number of aromatic residues nearby the chromophore retinal.

## Discussion

By measuring and analyzing the TAS and TFS for three mutants of the CRABPII-based rhodopsin mimic in acidic solution, we propose the excited-state dynamics of AT-PRSB in a low pH environment as shown in Fig 7, which is significantly different from the dynamics of the 15C-PRSB in the neutral solution as indicated in reference (**?**). In detail, immediately after being excited by the pump pulse, the system jumps to the FC state, followed by a fast relaxation (in about several hundreds of femtoseconds) to an intermediate state J. From the intermediate state J, the system further decays to a second intermediate state K and finally isomerize to the ground state of 15C-PRSB through the CI point, taking tens of picoseconds. In the end, the ground state of 15C-PRSB can relax back to the AT-PRSB state in a time scale of tens of nanoseconds or even longer according to our experiment. We can not determine the structures of the two intermediate states through the spectra obtained from current experiments. However, according to the hypothesis proposed in literature (17), we infer that the intermediate state J should be the excited state of 13C-15T-PSB while the intermediate state K should be the excited state of 13C-15C-PSB.

**Figure 7:**
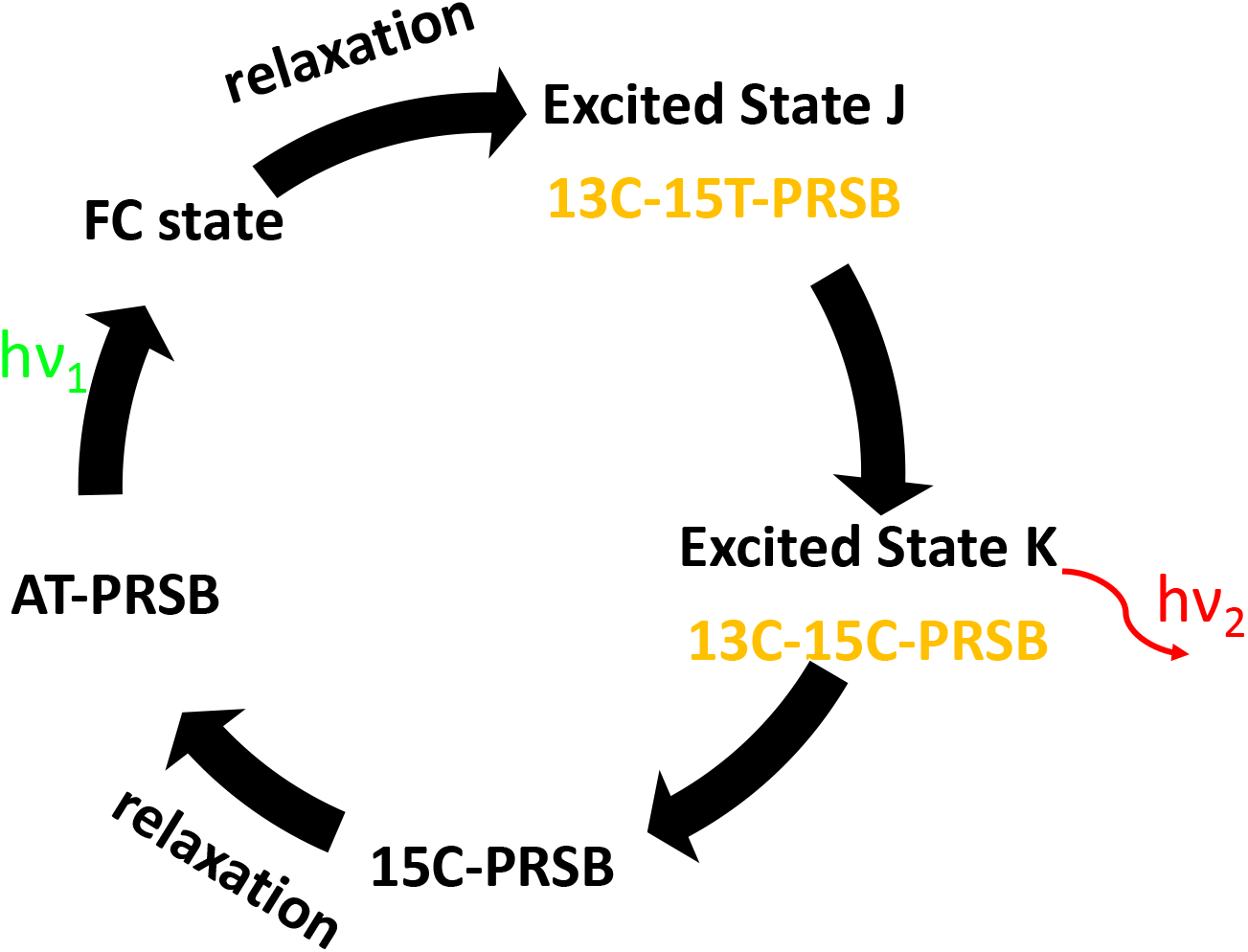
The proposed photoreaction cycle according to the experimental observations. The states colored in orange are not directly observed in the experiment but inferred according to the hypothesis proposed in reference(17).

Through the comparisons across three mutants, we find that mutant M9 has the longest wavelengths of maximum absorption and fluorescence emission, the longest lifetimes for the isomerization and the excited states. In contrast, M1 has the shortest wavelength and lifetime. Recalling the structures of the mutants as shown in Fig 1, we can see that the mutant M9 has three aromatic residues surrounding the *β*-ionone ring structure of the retinal, while M2 has two and M1 has only one. Based on the above observation, we infer that the aromatic residues nearby the retinal can not only lead to the red-shift of the spectra but also increase the stability of AT-PRSB, due to the *π* stacking effects between the benzyl-like structure of the sidechain in the aromatic residues and the *β*-ionone of the retinal, which is just similar to the case of designing the red fluorescence protein from the GFP using the triple-decker motif (23). Following this inference, one may further design new rhodopsin-like proteins that can emit fluorescence with even longer wavelength and lifetime, by mutating more non-aromatic residues near the *β*-ionone of retinal to aromatic ones. Such designable proteins could be highly useful in the fields of biosensing or bioimaging, as well as in some special cases when the traditional GFP is not applicable.(24).

## Conclusion

Synthetic rhodopsin mimics have great potential in the photosensitive devices due to their high stability and photosensitivity. To achieve this goal, understanding the photoreaction mechanism of the RSB in rhodopsin mimics is particularly important. In this article, we investigated the dynamics of the AT-PRSB in a low pH environment with three mutants of the CRABP-based rhodopsin mimic as examples. We measured the TAS and TFS for all the three mutants, from which the corresponding excited-states dynamics of the PRSB in acidic solution are obtained. We find that the three mutants share the same model on the photoreaction process. The different mutations in the CRABPII-based rhodopsin mimic only result in the shifts in the spectral wavelength and the lifetimes of the excited states. By correlating the structural properties with the observations in spectra, we propose a hypothesis that the aromatic residues nearby the *β*-ionone ring structure of the retinal could change the electron density along the reintal and increase the retinal’s stability, and hence lead to the red-shift of the spectra and increase in the lifetimes of the excited states, respectively. According to our current measurements, the mutant M9 already has a fluorescence emission in far red region with wavelength at 688nm. By performing further mutations on the neighboring residues near the *β*-ionone ring structure of the retinal, one could design a rhodopsin-like protein with infrared fluorescence, which can be particularly useful in the applications in biosensing or bioimaging in deeper tissues.

## Method

### Protein expression and purification

The mutants of the CRABPII-based rhodopsin mimic were prepared in a way similar to the protocols described in the literature(11; 25). The mutants of the CRABP-II protein were expressed from the pET-28a vector. The target gene was transformed into Escherichia coli BL21 (DE3) cells (300 ng of DNA, for 100 μL of cell solution) following standard protocols and the cells were grown on Luria-Bertani (LB)-agar plates supplemented with antibiotics (Kanamycin: 50μg/mL) at 37 ºC for 14 h. A single colony was used to inoculate 100 mL of LB medium containing 50 μg/mL Kanamycin and was grown at 37 ºC, while shaking overnight. The resulting culture was used to inoculate 1 L of LB containing 50 μg/mL Kanamycin and was grown at 37 ºC while shaking till OD_600_ reached 0.8. The expression was induced with addition of isopropyl-*β*-D-thiogalactopyranoside (IPTG, Gold Biotechnology, 0.5mM) and the culture was shaken at 37 ºC for 6 h. The cells were harvested by centrifugation (4000 rpm, 15 min, 4 ºC) and resuspended in Tris-Hcl buffer (25 mM Tris, 150mM NaCl,pH=8.0, 40 mL),disrupted by using a Ultra high pressure homogenizer.The solubilized membranes were isolated by high speed centrifugation (14000 rpm, 50 min, 4°C), and the supernatant was applied to a Ni-affinity column (HisTrap, GE Healthcare) at 4°C.After washed with Tris buffer, the mutants were eluted with a linear gradient of imidazole. The purified protein was concentrated by centrifugation using an Amicon Ultra filter. Samples were further purified using Akta pure L employing HiTrap Desalting anion (GE Health Sciences). The elution buffer was substituted with Tris-Hcl buffer using desalting columns. The obtained protein solution was titrated with HCl solution (6Mhcl) until the pH value reaches 3. The prepared samples for mutants M1,M2 and M9 are shown in Fig S4, where the colorful solutions indicate the RSB is well protonated.

### Steady-state spectral measurements

All-trans-retinal was purchased from Sigma-Aldrich and used as received without further purification. The samples were prepared according to the ratio of retinal to protein in the literature(11). Ultraviolet–visible absorption spectra in different mutants were recorded by a GE Ultrospec9000 spectrophotometer. Fluorescence spectra were recorded on a HORIBA UV-800C spectrophotometer. On the fluorescence spectral measurement, the sample was excited by the pulse at 560 nm, and the fluorescence emission was recorded in a range from 580 to 775nm.

### Transient absorption spectrum

The light source of the apparatus was a Ti:sapphire regenerative amplifier (35fs, 800 nm, 1 kHz). This pulse was split into two, which were used as the pump and probe pulses. Broad band probing white light was generated by focusing part of output of the laser to a CaF_2_ plate. The pump pulses generated 560nm excitation light with the energy 400nJ, through an optical parametric amplifier(TOPAS Prime, Spectra Physics). Polarization of the pump pulse was adjusted to the magic angle with respect to the horizontally polarized probe pulse. In the optical path, a precise electric time delay line was used to control the time delay between excitation light and detection light. The pump and probe pulses were focused into a 1 mm thick flow cell, in which the sample solution was circulated. The flow speed was adjusted so that each laser shot experienced a fresh solution. This prevents multiple excitation of molecules within the photocycle and accumulation of photoproducts. The absorbance of the samples was set to 0.4 at the 1 mm thick flow cell. Spectra of the probe pulse after passing through the sample and reference pulse were acquired with a spectrometer(AvaSpec-ULS2048CL-EVO, Avantes). The full width at half maximum (FWHM) of the instrumental response function was 150fs.

### Transient fluorescence spectrum

The 800 nm pulses from a Coherent Astrella regenerative amplifier (80 fs, 1 kHz), seeded by a Coherent Vitara-s oscillator (35 fs, 80 MHz), were used to pump an optical parametric amplifier (Coherent, OperA Solo) to generate an excitation pulse at 560 nm.The laser was focused on the sample with a 10× objective lens. Picosecond transient fluorescence spectra were recorded using a streak camera (Optronis). All measurements were performed at room temperature. The system has an ultimate temporal resolution of ~ 20ps.

### Global analysis on spectrum

Global analysis was carried out using the software package Glotaran (26). To find the proper number of compartments, we tried several different values to fit the spectra. If the number of compartments is large enough, the spectral can always be well fitted. However, the EADS of some compartments will be lack of structures and more like random noise. While keeping the 3D spectra in Fig S1 being well fitted, we gradually reduce the number of compartments until the EADS of all the compartments have clear structures.

### MD simulations

The MD simulations are carried out with software NAMD2(27) with the force field CHARMM36m (28) for the protein and ions with the modified TIP3P water model (29). The force field parameters for RSB are taken from the literature (30; 31). The system temperature was kept at 300 K and the system pressure was 1 atm. During the simulations, a 2 fs integrator time step was used with the bond vibrations constrained. Non-bonded forces are calculated every 2fs, and the full electrostatics forces are evaluated every 4fs with the particle mesh Ewald (PME) method(32). The cutoff for the non-bonded pair list is set to 14Å. The cutoff and switching distance for the non-bonded potential are set to 12Å and 10Å, respectively. The system was first minimized, then annealed to the target temperature from 60K by increasing the temperature at a rate of 1 K/ps, and then equilibrated for 1 ns. During these simulations, the protein backbone was constrained with a spring force constant of k = 2.0 kcal mol^−1^ Å^−2^. Finally, a production run without constraints was performed for 200 ns in total.

## Data availability

All data are contained within the article.

## Supporting information

This article contains supporting information.

## Acknowledgments

We acknowledge the Protein Preparation and Characterization Core Facility of Tsinghua University Branch of China National Center for Protein Sciences Beijing for providing the facility support.

## Author contributions

Q. Z. and X. P. conceptualization; Q. Z., G. L. and X. P. methodology; G. L., Y. H., S. P., J. M., J. W., J. W., S. Y. and Z. W. investigation; G. L. data curation; S. W., X. L.and Y. W. resources; G. L. and X. P. writing the draft; Q.Z. review and editing; G. L. and X. P. visualization; Q. Z. supervision; Q. Z. funding acquisition

## References

[1] Graham R Fleming, Gregory D Scholes, and Yuan-Chung Cheng. Quantum effects in biology. Procedia Chemistry, 3(1):38–57, 2011.

[2] Jennifer C Brookes. Quantum effects in biology: golden rule in enzymes, olfaction, photosynthesis and magnetodetection. Proceedings of the Royal Society A: Mathematical, Physical and Engineering Sciences, 473(2201):20160822, 2017.

[3] Neill Lambert, Yueh-Nan Chen, Yuan-Chung Cheng, Che-Ming Li, Guang-Yin Chen, and Franco Nori. Quantum biology. Nature Physics, 9(1):10–18, 2013.

[4] Steven O Smith. Structure and activation of the visual pigment rhodopsin. Annual review of biophysics, 39:309–328, 2010.

[5] Dieter Oesterhelt and Walther Stoeckenius. Rhodopsin-like protein from the purple membrane of halobacterium halobium. Nature new biology, 233(39):149–152, 1971.

[6] Oded Béja, Elena N Spudich, John L Spudich, Marion Leclerc, and Edward F DeLong. Proteorhodopsin phototrophy in the ocean. Nature, 411(6839):786–789, 2001.

[7] Akira Kawanabe and Hideki Kandori. Photoreactions and structural changes of anabaena sensory rhodopsin. Sensors, 9(12):9741–9804, 2009.

[8] Alina Pushkarev, Keiichi Inoue, Shirley Larom, José Flores-Uribe, Manish Singh, Masae Konno, Sahoko Tomida, Shota Ito, Ryoko Nakamura, Satoshi P Tsunoda, et al. A distinct abundant group of microbial rhodopsins discovered using functional metagenomics. Nature, 558(7711):595–599, 2018.

[9] Samer Gozem, Hoi Ling Luk, Igor Schapiro, and Massimo Olivucci. Theory and simulation of the ultrafast double-bond isomerization of biological chromophores. Chemical reviews, 117(22):13502–13565, 2017.

[10] Tetyana Berbasova, Meisam Nosrati, Chrysoula Vasileiou, Wenjing Wang, Kin Sing Stephen Lee, Ipek Yapici, James H. Geiger, and Babak Borhan. Rational design of a colorimetric ph sensor from a soluble retinoic acid chaperone. Journal of the American Chemical Society, 135(43):16111–16119, 2013.

[11] Meisam Nosrati, Tetyana Berbasova, Chrysoula Vasileiou, Babak Borhan, and James H Geiger. A photoisomerizing rhodopsin mimic observed at atomic resolution. Journal of the American Chemical Society, 138(28):8802–8808, 2016.

[12] Rachael M Crist, Chrysoula Vasileiou, Montserrat Rabago-Smith, James H Geiger, and Babak Borhan. Engineering a rhodopsin protein mimic. Journal of the American Chemical Society, 128(14):4522–4523, 2006.

[13] Alireza Ghanbarpour, Muath Nairat, Meisam Nosrati, Elizabeth M Santos, Chrysoula Vasileiou, Marcos Dantus, Babak Borhan, and James H Geiger. Mimicking microbial rhodopsin isomerization in a single crystal. Journal of the American Chemical Society, 141(4):1735– 1741, 2018.

[14] Wenjing Wang, Zahra Nossoni, Tetyana Berbasova, Camille T Watson, Ipek Yapici, Kin Sing Stephen Lee, Chrysoula Vasileiou, James H Geiger, and Babak Borhan. Tuning the electronic absorption of protein-embedded all-trans-retinal. Science, 338(6112):1340–1343, 2012.

[15] M Mark, R Konstantin, E Askat, H James, S Delmar, et al. Toward an understanding of the retinal chromophore in rhodopsin mimics. Journal of physical chemistry, 2013.

[16] Baptiste Demoulin, Margherita Maiuri, Tetyana Berbasova, James H Geiger, Babak Borhan, Marco Garavelli, Giulio Cerullo, and Ivan Rivalta. Control of protonated Schiff base excited state decay within visual protein mimics: a unified model for retinal chromophores. Chemistry–A European Journal, 27(66):16389–16400, 2021.

[17] Madushanka Manathunga, Adam J. Jenkins, Yoelvis Orozco-Gonzalez, Alireza Ghanbarpour, Babak Borhan, James H. Geiger, Delmar S. Larsen, and Massimo Olivucci. Computational and spectroscopic characterization of the photocycle of an artificial rhodopsin. The Journal of Physical Chemistry Letters, 11(11):4245–4252, 2020.

[18] Georgios N Stamatas, Michael Southall, and Nikiforos Kollias. In vivo monitoring of cutaneous edema using spectral imaging in the visible and near infrared. Journal of investigative dermatology, 126(8):1753–1760, 2006.

[19] Eric F Pettersen, Thomas D Goddard, Conrad C Huang, Gregory S Couch, Daniel M Greenblatt, Elaine C Meng, and Thomas E Ferrin. UCSF Chimera—a visualization system for exploratory research and analysis. Journal of computational chemistry, 25(13):1605– 1612, 2004.

[20] Maxim V Shapovalov and Roland L Dunbrack Jr. A smoothed backbone-dependent rotamer library for proteins derived from adaptive kernel density estimates and regressions. Structure, 19(6):844–858, 2011.

[21] Ivo HM van Stokkum, Delmar S Larsen, and Rienk Van Grondelle. Global and target analysis of time-resolved spectra. Biochimica et Biophysica Acta (BBA)-Bioenergetics, 1657(2-3):82–104, 2004.

[22] AV Deshpande, A Beidoun, Alfons Penzkofer, and G Wagenblast. Absorption and emission spectroscopic investigation of cyanovinyldiethylaniline dye vapors. Chemical physics, 142(1):123–131, 1990.

[23] Computational challenges in modeling of representative bioimaging proteins: GFP-like proteins, flavoproteins, and phytochromes, author=Nemukhin, Alexander V and Grigorenko, Bella L and Khrenova, Maria G and Krylov, Anna I. The Journal of Physical Chemistry B, 123(29):6133–6149, 2019.

[24] Hannah E Chia, E Neil G Marsh, and Julie S Biteen. Extending fluorescence microscopy into anaerobic environments. Current opinion in chemical biology, 51:98–104, 2019.

[25] Rachael M. Crist, Chrysoula Vasileiou, Montserrat Rabago-Smith, James H. Geiger, and Babak Borhan. Engineering a rhodopsin protein mimic. Journal of the American Chemical Society, 128(14):4522–4523, 2006. PMID: 16594659.

[26] Joris J. Snellenburg, Sergey Laptenok, Ralf Seger, Katharine M. Mullen, and Ivo H. M. van Stokkum. Glotaran: A Java-Based Graphical User Interface for the R Package TIMP. Journal of Statistical Software, 49(3):1–22, 2012.

[27] James C Phillips, Rosemary Braun, Wei Wang, James Gumbart, Emad Tajkhorshid, Elizabeth Villa, Christophe Chipot, Robert D Skeel, Laxmikant Kale, and Klaus Schulten. Scalable molecular dynamics with NAMD. Journal of computational chemistry, 26(16):1781–1802, 2005.

[28] Jing Huang, Sarah Rauscher, Grzegorz Nawrocki, Ting Ran, Michael Feig, Bert L. de Groot, Helmut Grubmüller, and Alexander D. MacKerell Jr. CHARMM36m: an improved force field for folded and intrinsically disordered proteins. Nature Methods, 14(1):71–73, January 2017.

[29] William L Jorgensen, Jayaraman Chandrasekhar, Jeffry D Madura, Roger W Impey, and Michael L Klein. Comparison of simple potential functions for simulating liquid water. The Journal of chemical physics, 79(2):926–935, 1983.

[30] Shigehiko Hayashi and Iwao Ohmine. Proton transfer in bacteriorhodopsin: structure, excitation, IR spectra, and potential energy surface analyses by an ab initio QM/MM method. The Journal of Physical Chemistry B, 104(45):10678–10691, 2000.

[31] Shigehiko Hayashi, Emad Tajkhorshid, Eva Pebay-Peyroula, Antoine Royant, Ehud M Landau, Javier Navarro, and Klaus Schulten. Structural determinants of spectral tuning in retinal proteins bacteriorhodopsin vs sensory rhodopsin II. The Journal of Physical Chemistry B, 105(41):10124–10131, 2001.

[32] Tom Darden, Darrin York, and Lee Pedersen. Particle mesh Ewald: An Nlog (N) method for Ewald sums in large systems. The Journal of chemical physics, 98(12):10089–10092, 1993.

